# Cochaperones enable Hsp70 to fold proteins like a Maxwell’s demon

**DOI:** 10.1101/283143

**Authors:** Huafeng Xu

## Abstract

The heat shock protein 70 (Hsp70) chaperones, vital to the proper folding of proteins inside cells, consume ATP and require cochaperones in assisting protein folding. It is unclear whether Hsp70 can utilize the free energy from ATP hydrolysis to fold a protein into a native state that is thermodynamically unstable in the chaperone-free equilibrium. Here we present a model of Hsp70-mediated protein folding, which predicts that Hsp70, as a result of differential stimulation of ATP hydrolysis by its Hsp40 cochaperone, dissociates faster from a substrate in fold competent conformations than from one in misfolding-prone conformations, thus elevating the native concentration above and suppressing the misfolded concentration below their respective equilibrium values. Previous models would not make or imply these predictions, which are experimentally testable. Our model quantitatively reproduces experimental refolding kinetics, predicts how modulations of the Hsp70/Hsp40 chaperone system affect protein folding, and suggests new approaches to regulating cellular protein quality.

## Introduction

The discovery of chaperones and their roles in assisting protein folding amended the long-held view that proteins spontaneously fold into their native structures^1–3^. Large, multi-domain proteins may take many hours to fold, or fail to fold properly altogether on their own^2,4^. ATP-consuming chaperones—including Hsp70s—provide critical assistance in the in vivo folding and the biological functions of broad sets of substrate proteins^3^. Extensive experimental studies have firmly established that the Hsp70 chaperones can greatly accelerate the folding of their substrate proteins^5,6^. Despite tremendous progress in the mechanistic studies of the Hsp70 chaperones^2,7^, including the development of theoretical models^8–10^, it remains unclear why ATP consumption is indispensable to these chaperones; enzymes do not need to consume free energy in catalyzing chemical reactions. Recently it was demonstrated that chaperones such as GroEL and Hsp70 depend on continuous ATP hydrolysis to maintain a protein in a native state that is thermodynamically unstable^11^, but it is unknown how Hsp70 can utilize the ATP free energy to alter the folding equilibrium. In addition, Hsp70s require Hsp40—also known as J proteins^12^— cochaperones in assisting protein folding. It is yet unexplained why cochaperones are absolutely necessary.

The Hsp70 chaperones, such as the bacterial DnaK, consist of an N-terminal nucleotide binding domain (NBD) and a C-terminal substrate binding domain (SBD). The Hsp70 SBD adopts an open conformation when its NBD is ATP-bound (we call the Hsp70 to be in the ATP-state), which allows the substrate to bind and unbind at high rates, whereas when the NBD is ADP-bound (ADP-state), the SBD changes to a closed conformation, rendering both binding and unbinding orders-of-magnitude slower^6,13^,^14^. Hsp70s have low basal ATP hydrolysis activities^15^. The Hsp40 cochaperones, such as the bacterial DnaJ, can drastically stimulate the ATPase activity of Hsp70 using their N-terminal J domain (JD)^16,17^, shared by all Hsp40s (hence the name J proteins)^12^. Hsp40s also have a C-terminal domain (CTD) that can bind to denatured proteins^18,19^. Both Hsp70 and Hsp40 recognize exposed hydrophobic sites^7,20^,^21^. As a result, they can distinguish different protein conformations using the corresponding difference in the exposed hydrophobic sites. For example, Hsp70 has been shown to bind to both unfolded and partially folded, near native protein structures, but not to native structures^22,23^. Hsp40 and Hsp70 may simultaneously bind to different segments of the same substrate molecule, and the consequent spatial proximity then facilitates the J domain binding to Hsp70 and accelerating its ATP hydrolysis^24–26^. Following ATP hydrolysis, the chaperone returns from the ADP-state to the ATP-state through nucleotide exchange, which is often catalyzed by nucleotide exchange factors (NEF) such as the bacterial GrpE^27,28^.

A protein may fold to its native state, *N,* or go into a misfolded/aggregated—we will use these two terms interchangeably—state, *M* (Fig. 1a). It is unknown whether Hsp70 can use the free energy from ATP hydrolysis to drive its substrate toward the native state such that *f*_*N*_ / *f*_*M*_ > *f*_*N,eq*_ / *f*_*M,eq*_, where *f*_*S*_ is the fraction of the substrate in state *S* at the steady state of Hsp70-mediated folding, and *f*_*S,eq*_ is the corresponding fraction at the folding equilibrium in the absence of the chaperone. Previous models^9,29^ mostly considered the chaperone as an unfoldase/holdase—which need not consume free energy—that pulls the substrate out of the misfolded state and holds it in an unfolded state. It was proposed that the free energy from ATP hydrolysis was used to achieve ultra-affinity in substrate binding^8,30^. As an unfoldase/holdase, Hsp70 would also pull the substrate out of the native state into the unfolded state; unless Hsp70 has a higher affinity for the native substrate than for the misfolded substrate, these models would predict *f*_*N*_ / *f*_*M*__*N,eq*_ / *f*_*M,eq*_.

Here we propose a model of Hsp70-mediated protein folding, in which Hsp70 and Hsp40 together constitute a molecular machine that uses the free energy from ATP hydrolysis to actively drive a protein toward its native state, so that *f*_*N*_ / *f*_*M*_ > *f*_*N,eq*_ / *f*_*M,eq*_. It suggests that without Hsp40, Hsp70 alone cannot change the ratio *f*_*N*_ / *f*_*M*_ from the equilibrium value *f*_*N,eq*_ / *f*_*M,eq*_. Our model thus answers the question why Hsp70 requires both the Hsp40 cochaperones and ATP consumption in assisting protein folding. Our model explains the puzzling non-monotonic dependency of folding efficiency on the chaperone and cochaperone concentrations. It makes quantitative predictions on how protein folding is affected by modulations of the chaperone environment, including changes in the ATPase activity or the nucleotide exchange rate of Hsp70. These predictions may be readily tested by experiments, and inform rational approaches to manipulating chaperone-mediated protein folding.

**Figure 1.**
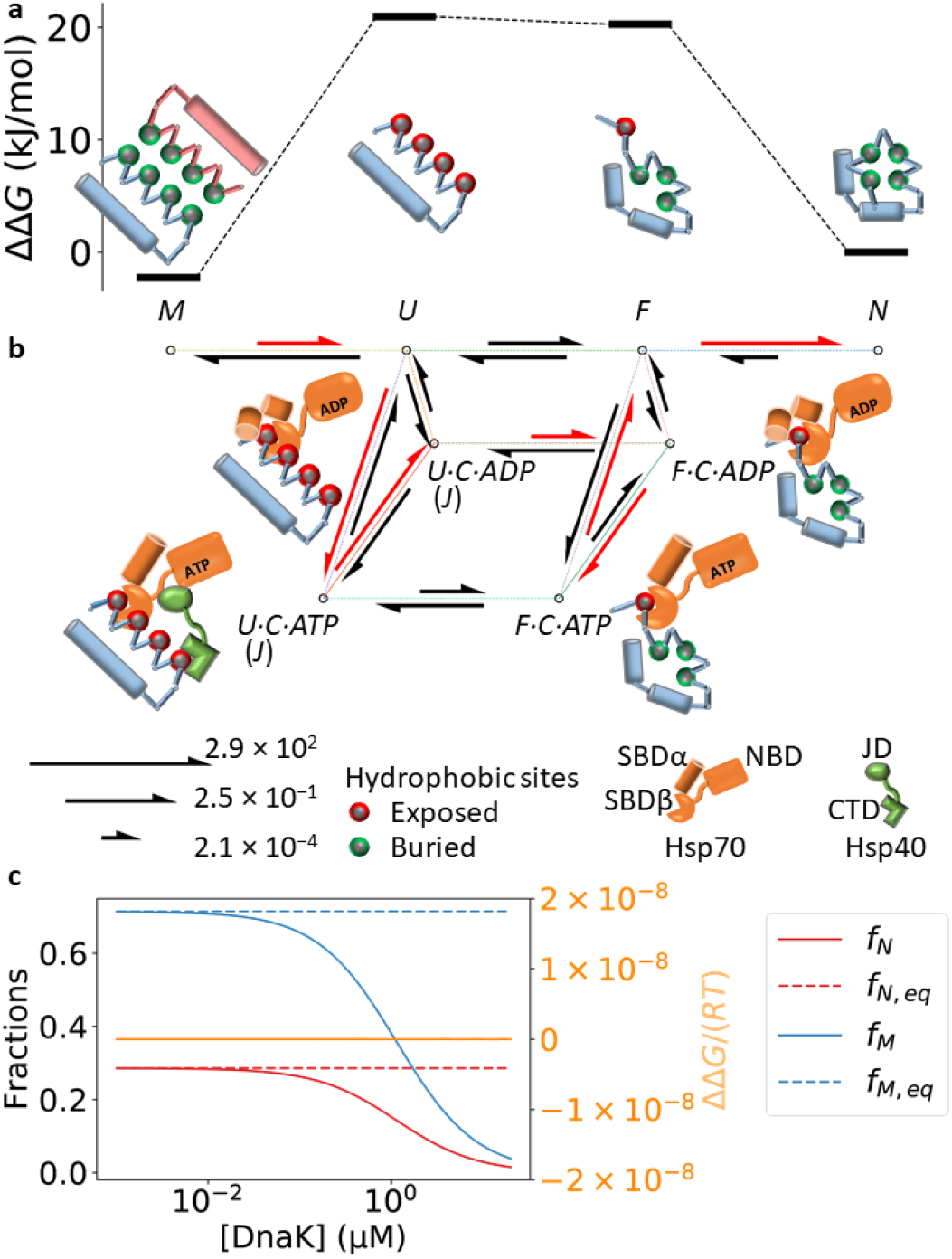
Our model of Hsp70/Hsp40/NEF-mediated protein folding. **a** Conformational states in protein folding, and their relative free energies in the absence of chaperones. We assume that there are four conformational states: *M*isfolded/aggregated, *U*nfolded and aggregation-prone, *F*old-competent, and *N*ative. The pink and blue chains in the *M* state may correspond to different molecules (aggregated) or to different domains in the same molecule (misfolded). The high free energy barriers associated with the intermediate states *U* and *F* slow down the refolding of misfolded proteins. The exemplary free energies are derived from the kinetic parameters fit to the refolding experiments of luciferase at 25 °C (Table 2). **b** The transitions between the microscopic states in the chaperone-mediated folding pathway. *S·C·X* represents the complex between the substrate in the conformational state *S* (*= U, F)* and the Hsp70 chaperone (denoted as *C*) bound to nucleotide *X (= ATP, ADP).* The transitions between *S* and *S·C·X* correspond to the chaperone binding to and unbinding from the substrate. The transition of *S·C·ATP* to *S·C·ADP* corresponds to ATP hydrolysis, and its reverse, nucleotide exchange. Hsp70 binding stabilizes the substrate in the intermediate states, thus catalyzing the folding reaction. Hsp40 (*J*) can form a ternary complex with the substrate and Hsp70—thus stimulating ATP hydrolysis—if the substrate is in the *U* state, but not if the substrate is in the *F* state. Differential ATP hydrolysis by Hsp70 bound to the substrate in the *U* and *F* states drives the refolding through the pathway highlighted in red. The lengths of the reaction arrows are linear with respect to the logarithms of the exemplary rate constants (in 1/s) for the DnaK/DnaJ/GrpE-mediated refolding of luciferase at 25 °C (Table 1, 2). **c** Without cochaperones, Hsp70 cannot alter the balance between folding and misfolding. DnaK binding to the intermediate states decreases both the native (red) and misfolded (blue) populations, but the ratio between the two remains unchanged from its equilibrium value: ΔΔ*G* = 0 (orange, right y-axis). Here, we have taken 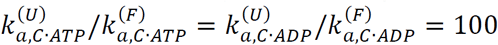 to show that differential binding of Hsp70 to the substrate in different conformational states does not alter the folding/misfolding balance.

## Results

In our model (detailed in *Methods*), we consider two additional conformational states—besides the *M* and *N* states—of a protein: the unfolded and aggregation-prone state, *U,* and the fold-competent state, *F.* A protein in the *F* state is unfolded but poised to fold into the native state (Fig. 1a, b). Such intermediate states of folding have been observed experimentally^4^. Conformational transitions can occur between *M* and *U,* between *U* and *F,* and between *F* and *N* (Fig. 1b). We assume that a protein in the *U* state has more exposed hydrophobic sites than in the *F* state—which is consistent with the experimental observations^4^—and a protein in the *M* and *N* states has nearly zero such sites, as both folding and aggregation (including oligomerization) bury the protein’s hydrophobic sites (Fig. 1a).

Key to our model is the assumption that Hsp40, like Hsp70, has different affinities for the substrate in different conformations^20^, favoring conformations with more exposed hydrophobic sites. Thus Hsp70 and Hsp40 can bind to a substrate molecule in the *U* and *F* states—with higher affinities for the *U* state than for the *F* state—but not to one in the *M* and *N* states (Fig. 1b). A substrate in the *U* state is more likely to be Hsp40-bound than one in the *F* state. As a result, an Hsp70 molecule bound to a substrate molecule in the *U* state will on average have substantially higher ATP hydrolysis rate—because of the more probable *cis* stimulation by an Hsp40 molecule bound to the same substrate molecule—than if it is bound to a substrate molecule in the *F* state. If the nucleotide exchange rate is between these two hydrolysis rates, an Hsp70 bound to a substrate in the *U* state will be driven toward the ADP-state, where it slowly dissociates from the substrate, while an Hsp70 bound to a substrate in the *F* state will be driven toward the ATP-state, where it rapidly dissociates from the substrate. Acting like a Maxwell’s demon^31^, Hsp70 releases the fold-competent substrate but retains the aggregation-prone substrate, driving the folding along the reaction path of *M* → *U* → *U · C · ATP* → *U · C · ADP* → *F · C · ADP* → *F C ATP → F → N* (*S · C · X* represents the complex between a substrate in conformation *S* and the chaperone *C* bound to nucleotide *X = ATP, ADP*) (Fig. 1b). One ATP molecule is consumed in this reaction path and the free energy is used to compel the substrate into the native state.

The extent to which Hsp70 biases protein folding can be quantified by the excess free energy: ΔΔ*G ≡ R T* (ln(*f*_*N*_ / *f*_*M*_) - ln(*f*_*N,eq*_ / *f*_*M,eq*_)), where *R* is the gas constant and *T* the temperature. A positive excess free energy requires not that more chaperones bind to the substrate in the *U* state than to the substrate in the *F* state, which is true and reflected in previous models, but that an individual chaperone molecule, when bound to a substrate, resides longer on it if the substrate is in the *U* state than if it is in the *F* state. For this, Hsp70 needs both ATP consumption and a cochaperone: it can be shown algebraically (see *Methods*) and numerically (Fig. 1c) that without cochaperones, ΔΔ*G* = 0. These predictions are consistent with the results from the single molecule experiment of DnaK-mediated refolding^32^, where DnaK alone in the presence of ATP was unable to alter the ratio of the misfolded and folded fractions.

We applied our model to the analysis of DnaK/DnaJ/GrpE-mediated refolding of luciferase^5^ and its variant LucDHis6^29^. Most of the relevant kinetic parameters for this bacterial Hsp70 system have been carefully determined experimentally^33^ (Table 1). Our model quantitatively reproduces the experimentally observed refolding kinetics under various conditions, capturing the slow spontaneous refolding and denaturation of luciferase, the acceleration of refolding with chaperone assistance, and the necessity of GrpE (Fig. 2a, b). The refolding speed and yield reach a maximum at the DnaK concentration of 1μM, which is captured by our model (Fig. 2b, c). The intermediate conformations *U* and *F* in our model may correspond to the experimentally identified intermediate conformations *I*_2_ and *I*_1_ of luciferase^4^: the free energy difference between *N* and *F* at 25 °C, according to the fitted parameters, is 20 kJ/mol, close to the experimental value of 15 kJ/mol between *N* and *I*_1_, measured at 10 °C. Consistent with previous experimental observations^29^, our model suggests that the Hsp70-accelerated refolding proceeds in two steps: 1) rapid unfolding of the misfolded substrate, stabilized by the ADP-bound DnaK, followed by 2) slow conversion to the native state (Fig. 3a).

**Table 1.** The experimental kinetic parameters for the DnaK/DnaJ/GrpE chaperone system. All the parameters are determined at the temperature of 25 °C, except for the rate of conformational change of DnaK from closed to open conformations, which was measured at 15 °C, 25 °C, and 35 °C, yielding the parameters 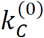 and *E*_*a*_ from a fitting of the Arrhenius equation to the measured rates at different temperatures.

| Reaction | Parameter | Rate | Reference |
| --- | --- | --- | --- |
| $S + C \cdot ATP \rightleftharpoons S \cdot C \cdot ATP$ | $k_{a,C \cdot ATP}^{(0)}$ | $1.28 \times 10^6 \text{ M}^{-1} \cdot \text{s}^{-1}$ | 6 |
| | $k_{d,C \cdot ATP}$ | $2.31 \text{ s}^{-1}$ | 6 |
| $S + C \cdot ADP \rightleftharpoons S \cdot C \cdot ADP$ | $k_{a,C \cdot ADP}^{(0)}$ | $1.17 \times 10^4 \text{ M}^{-1} \cdot \text{s}^{-1}$ | 6 |
| | $k_{d,C \cdot ADP}$ | $9.1 \times 10^{-4} \text{ s}^{-1}$ | 6 |
| $J + C \cdot ATP \rightleftharpoons J \cdot C \cdot ATP$ | $k_{a,J \cdot C}$ | $4.88 \times 10^4 \text{ M}^{-1} \cdot \text{s}^{-1}$ | 15<sup>a</sup> |
| | $k_{d,J \cdot C}$ | $1.6 \times 10^{-3} \text{ s}^{-1}$ | 46 |
| $S \cdot C \cdot ATP \rightarrow S \cdot C \cdot ADP + Pi$ | $k_h^{(0)}$ | $0.01 \text{ s}^{-1}$ | 15<sup>b</sup> |
| $J \cdot S \cdot C \cdot ATP \rightarrow J \cdot S \cdot C \cdot ADP + Pi$ | $k_h^{(J)}$ | $1.8 \text{ s}^{-1}$<sup>c</sup> | 15 |
| $C \cdot ADP \rightarrow C + ADP$ | $k_{d,ADP}$ | $0.006 \text{ s}^{-1}$ | 47 |
| $E + C \cdot ADP \rightleftharpoons E \cdot C \cdot ADP$ | $k_{a,E \cdot C}$ | $7.85 \times 10^6 \text{ M}^{-1} \cdot \text{s}^{-1}$ | 27<sup>d</sup> |
| | $k_{d,E \cdot C}$ | $30 \text{ s}^{-1}$ | 27 |
| $C_{\text{closed}} \cdot ATP \rightarrow C_{\text{open}} \cdot ATP$ | $k_C^{(0)}$ | $1.1 \times 10^{23} \text{ s}^{-1}$ | 45<sup>e</sup> |
| | $E_a/R$ | $17320 \text{ K}$ | 45 |

**Figure 2.**
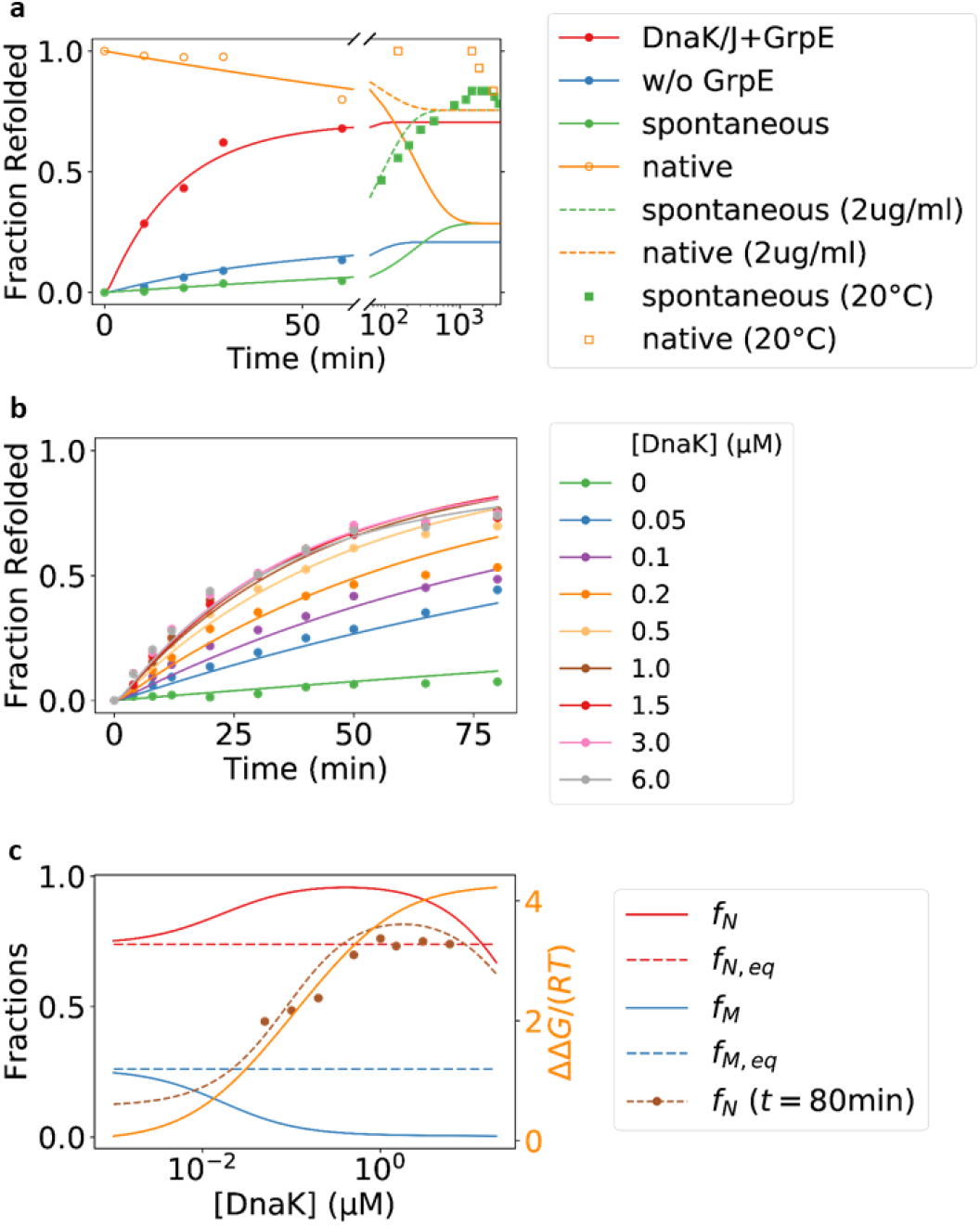
Our model is in good agreement with previous experimental studies of DnaK/DnaJ/GrpE-mediated refolding, and it predicts that DnaK and DnaJ cooperate to utilize the free energy of ATP hydrolysis to drive luciferase toward the native state. **a** The refolding of denatured luciferase under various conditions. The predictions of our model are shown in lines, whereas the experimental data are shown as filled circles^5^. The dashed lines show the spontaneous refolding and denaturation at the much lower substrate concentration of 0.032 μM (2 μg/ml), as in the corresponding experiments (empty circles and squares)^4^. The experiments of spontaneous refolding and denaturation were performed at 20 °C, lower than the temperature of 25 °C at which most of the kinetic parameters were obtained. In modeling the refolding, we assume that initially all the protein is in the misfolded/aggregated state, i.e., *f*_*M*_ (*t* = 0) = 1; in modeling the denaturation, we assume that initially all the protein is in the native state, i.e., *f*_*N*_ (*t* = 0) = 1. **b** The refolding of a luciferase mutant, LucDHis6^29^, in the presence of DnaK at various concentrations. Although the experiments were performed at the slightly lower temperature of 22°C, the predictions of our model using the kinetic parameters derived for 25°C are nevertheless in quantitative agreement with the experimental data. **c** The native fraction (*f*_*N*_ = [*N*] / [*S*], red) and the misfolded fraction (*f*_*M*_ = [*M*] / [*S*], blue) of LucDHis6 at the steady state of DnaK-mediated refolding at various DnaK concentrations. The corresponding fractions in the chaperone-free folding equilibrium, *f*_*N,eq*_ and *f*_*M,eq*_, are shown as dashed lines. The unitless excess free energy ∆*∆G* / (*RT*) is shown in orange (right y-axis). The fractions after 80 min of refolding, starting from misfolded LucDHis6, are shown in brown. Our model is in good agreement with the experimental data (filled circles), and it suggests that the refolding is still incomplete even after 80 min. The fitting parameters in our model are given in Table 2, and the conditions of the experiments considered in this paper are summarized in Table 3.

At the steady state, the reactive flux along the ATP-driven cycle *U → U C ATP → U C · ADP* → *F · C · ADP* → *F · C · ATP* → *F* (*→ U*) (Fig. 3b) keeps the protein folding out of equilibrium, elevating the native population above and suppressing the misfolded population below their respective equilibrium values (Fig. 2c). The excess free energy at the steady state always increases with increasing DnaK concentrations, but the native population reaches a maximum and then decreases (Fig. 2c), because at high DnaK concentrations, the substrate is trapped in the DnaK-bound state and thus prevented from folding into the native state.

**Table 2.** The model parameters fit to the refolding experiments.

| Reaction | Parameter | Luciferase |  | LucDHis6 |
| --- | --- | --- | --- | --- |
|  |  | 25 °C | 30 °C | 22 °C |
| $M \rightleftharpoons U$ | $k_{M \rightarrow U} (s^{-1})^a$ | 0.02 | 0.02 | 0.02 |
| | $k_{U \rightarrow M} (M^{-1} \cdot s^{-1})$ | $9.4 \times 10^8$ | $3.4 \times 10^9$ | $6.4 \times 10^7$ |
| $U \rightleftharpoons F$ | $k_{U \rightarrow F} (s^{-1})$ | 0.23 | 0.88 | 0.26 |
| | $k_{F \rightarrow U} (s^{-1})$ | 0.18 | 0.13 | 0.042 |
| $F \rightleftharpoons N$ | $k_{F \rightarrow N} (s^{-1})^b$ | 5 | 5 | 5 |
| | $k_{N \rightarrow F} (10^{-3} \cdot s^{-1})$ | 1.4 | 20 | 1.1 |
| $S + C \cdot X \rightleftharpoons S \cdot C \cdot X$ | $n_c^{(U)}$ | 9.1 | 6.2 | 1.9 |
| | $n_c^{(U)}/n_c^{(F)}^c$ | 20 | 20 | 20 |
| $S + J \rightleftharpoons S \cdot J$ | $K_{A,J}^{(U)} (10^6 \cdot M^{-1})$ | 13 | 22 | 54 |
| $(J + C) \cdot S \rightleftharpoons (J \cdot C) \cdot S$ | $[J]_{\text{eff}} (M)^d$ | $5 \times 10^{-3}$ | $5 \times 10^{-3}$ | $5 \times 10^{-3}$ |

- The predictions of our model depend on the equilibrium constant $K_{M \rightleftharpoons U} = k_{M \rightarrow U}/k_{U \rightarrow M}$ , but are degenerate with respect to the individual rates. We thus fix an arbitrary but plausible value for $k_{M \rightarrow U}$ and fit $k_{U \rightarrow M}$ to the experimental refolding data. - For the similar reason as above, we fix $k_{F \rightarrow N}$ and fit $k_{N \rightarrow F}$ to the refolding data. - There should be more accessible DnaK binding sites in the $U$ state than in the $F$ state. Here, we arbitrarily set the ratio between the two. Although the values of the other fitting parameters will change accordingly, the quality of the fit and the predictions of our model are insensitive to this ratio (at least for values between 1 and 100). - We assume that the effective distance, $L$ , between the Hsp70 molecule and the J domain of the Hsp40 molecule bound to the same substrate molecule is $L = 4.3$ nm, the effective concentration is then $[J]_{\text{eff}} = N_A^{-1}/(L^3 4\pi/3) = 5 \times 10^{-3} M$ . The quality of the fit and the predictions of our model are insensitive to this parameter.

The model parameters fit to the refolding experiments. of LucDHis6 (Fig. 4). In the initial minutes of refolding, approximately 150 ATP molecules are consumed to refold one LucDHis6 (Fig. 4a), which is reasonably close to the experimental result of ~50 ATP molecules consumed per refolded LucDHis6 when the stoichiometry of DnaK:LucDHis6 is 1:1, significantly higher than the experimental number of ~5 when LucDHis6 is in excess of DnaK, and significantly lower than the estimates of > 1000 for many other substrates in other experiments^29^,^34–36^. The discrepancy between the model and the experimental results may be partially attributable to the approximations in our model and the inaccuracies in the input kinetic parameters. ATP hydrolysis continues at the steady state and the free energy is utilized to promote the native state and suppress the misfolded state (Fig. 4b, c). As [DnaK] exceeds 1 µM, the ATP consumption rate increases rapidly without commensurate increase in the excess free energy. Our analysis thus suggests that DnaK may be most free energy efficient at maintaining protein folding out-of-equilibrium when its concentration is in the sub-micromolar range, a prediction that may be tested experimentally.

Our model suggests that Hsp70 can keep a protein folded even if it thermodynamically favors aggregation. The chaperone is thus able to play a critical role in maintaining protein conformations, not just in the folding of nascent chains^37^. Higher DnaK concentrations are required to suppress aggregation at increasing substrate concentrations (Fig. 5a) or at decreasing substrate stabilities (Fig. 5b). This may explain how cells that overexpress DnaK can tolerate higher numbers of mutations in the chaperone’s substrates^38^. Because the excess free energy plateaus at high chaperone concentrations (Fig. 2c), our results imply a limit on the chaperones’capacity to prevent aggregation, in that there exists a threshold of aggregation tendency (Fig. 5a, b, the black arrows) above which the chaperone can no longer maintain high levels of native concentrations and prevent aggregation at the same time.

**Figure 3.**
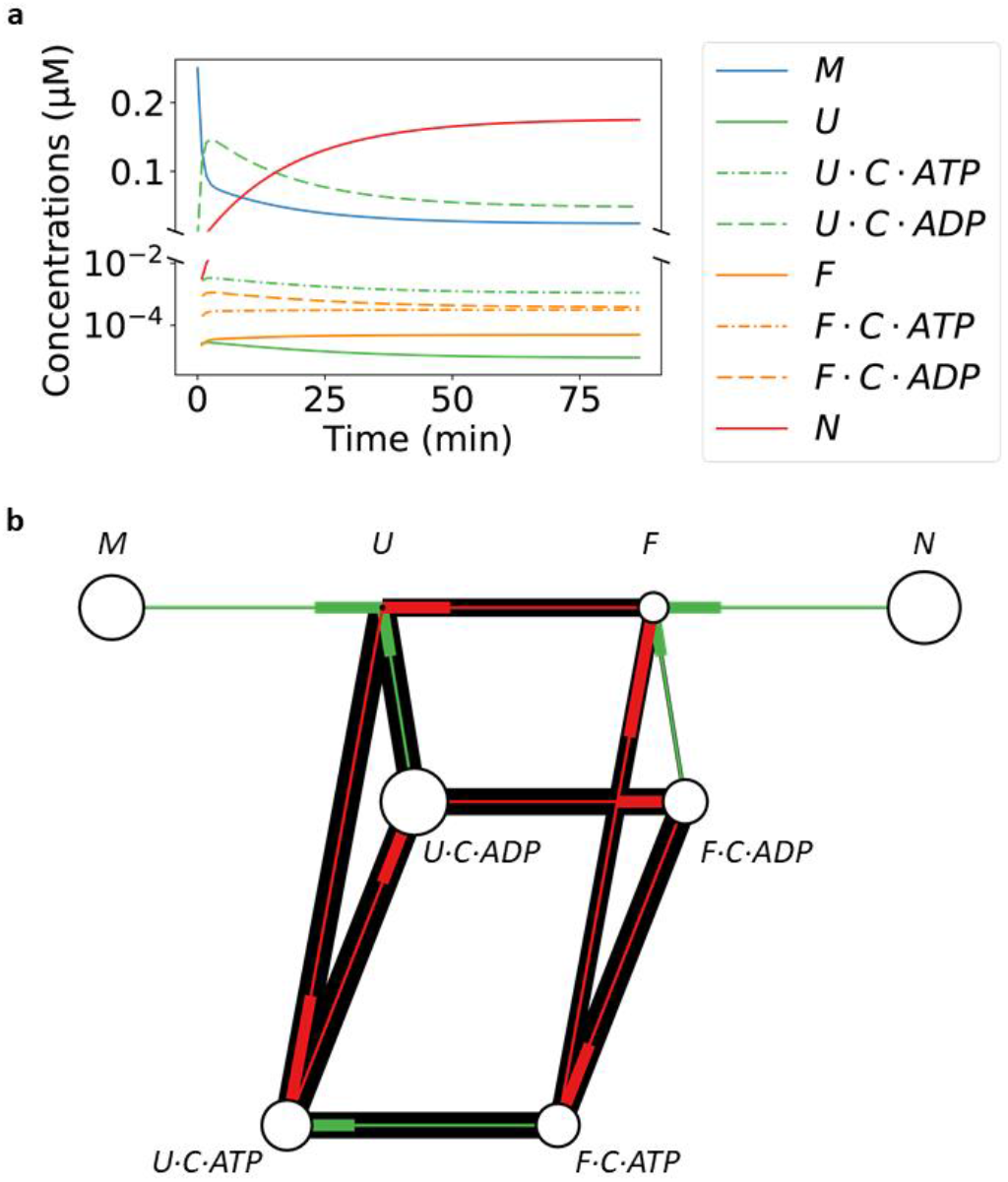
The mechanism by which the Hsp70 chaperone system accelerates refolding and maintains the folding out of equilibrium. **a** The model prediction of the concentrations of different molecular species in DnaK/DnaJ/GrpE-mediated refolding of denatured luciferase. The substrate is primarily held by ADP-bound chaperone in the *U* state before slowly moving into the native state. **b** The reactive flux at the steady state. The thickness of each line is linear with respect to the logarithm of the absolute value of the reactive flux, and the thicker end of the thin, center line indicates the destination of the flux. The size of the circle at each node is linear with respect to the logarithm of the steady state fraction of the corresponding molecular species. The ATP-driven reaction cycle is highlighted in red.

Our model suggests that Hsp70 only drives the folding of proteins with sufficiently slow conversion between *U* and *F* states (Fig. 5c, d, e), implying that Hsp70 substrates tend to be slow refolding proteins (Fig. 5d). If the conversion between *U* and *F* is too fast, the chaperone diminishes, rather than increases, the native fraction in comparison to the chaperone-free equilibrium. As the conversion slows, the chaperone drives the steady state native fraction higher, but at the price of longer refolding time (Fig. 5e), a trade-off reminiscent of that between speed and specificity in the kinetic proofreading mechanism^39,40^, where the expenditure of free energy (such as from ATP or GTP consumptions) is utilized to increase the specificity of chemical reactions.

Our model explains the observation that folding is less efficient at both low and high DnaJ concentrations^15^ (Fig. 6a). At low DnaJ concentrations, ATP hydrolysis is slow, and nucleotide exchange drives DnaK toward the ATP-state, in which it dissociates from the substrate rapidly and thus unable to prevent aggregation. At high DnaJ concentrations, a large fraction of the substrate in the *U* state is bound to DnaJ. These DnaJ-bound substrate molecules are trapped in the *U* state, unable to progress toward the *F* state, resulting in diminished folding.

**Figure 4.**
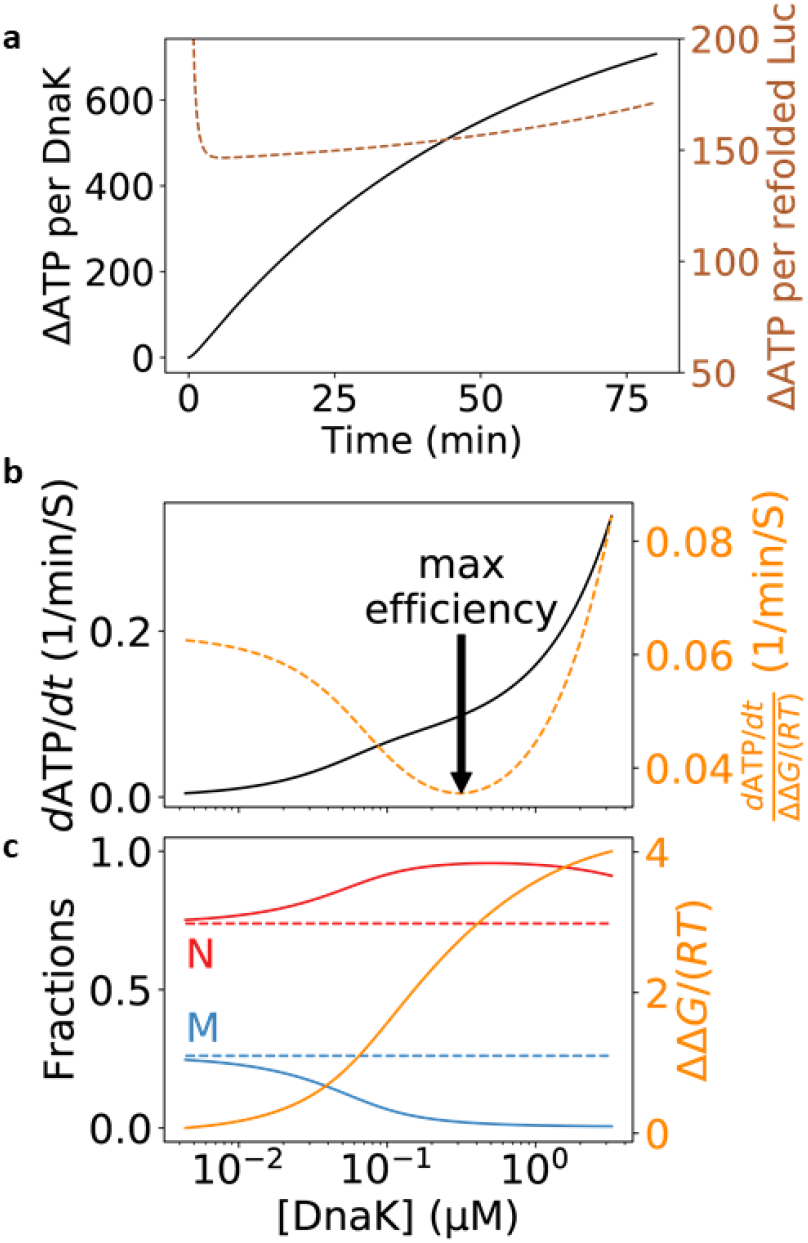
Free energy consumption in DnaK/DnaJ/GrpE-mediated folding of LucDHis6. **a** ATP consumption in the course of refolding of denatured LucDHis6. The instantaneous rate of ATP consumption is given by 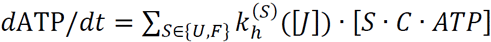, and the number of hydrolyzed ATP per molecule of refolded substrate can be estimated by dividing the cumulative consumption, 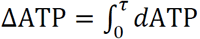, by the number of refolded substrate molecules after time *τ*. Here, the DnaK concentration is 0.5 μM, and the other kinetic parameters are given in Table 1 and Table 2. The black curve shows the number of ATP molecules hydrolyzed per DnaK molecule after the given time. The brown curve shows the number of ATP molecules consumed per one molecule of refolded LucDHis6 (right y-axis) up to the given time, which increases to infinity at the steady state because no additional LucDHis6 is refolded, yet ATP hydrolysis continues. **b** and **c** ATP hydrolysis at the steady state. The ATP consumption rate per substrate, at various DnaK concentrations, is shown as the black curve in panel b, and the corresponding native (red) and misfolded (blue) fractions at the steady state are shown as solid lines in panel c. The native fraction is above and the misfolded fraction is below their respective equilibrium values (dashed flat lines). The excess free energy ΔΔ*G* / (*RT*) is shown in orange (right y-axis) in panel c. We can measure the chaperones’ free energy efficiency in maintaining the non-equilibrium by the ratio of the ATP consumption rate to the excess free energy at the steady state (orange, right y-axis, in panel b). The arrow indicates the DnaK concentration at which the chaperones utilize the least amount of ATP per unit of excess free energy.

Our model also explains the observation that folding decreases at both low and high GrpE concentrations^27^ (Fig. 6b). For the chaperone to effectively assist folding, nucleotide exchange should be much slower than ATP hydrolysis when the chaperone binds to a substrate in the *U* state, but much faster than ATP hydrolysis when it binds to a substrate in the *F* state, so that the chaperone is driven toward the ADP-state in the former case, and toward the ATP-state in the latter case (Fig. 1b). At low GrpE concentrations, nucleotide exchange is slow, leaving DnaK bound to the substrate in the *F* state predominantly in the ADP-state—as reflected by the low population of *F · C · ATP* (Fig. 6b), slowing its dissociation from the substrate and thus preventing the latter from folding to the native state. At high GrpE concentrations, nucleotide exchange is fast, and DnaK is driven into the ATP-state and does not stay bound to the substrate in the *U* state long enough—as reflected by the decreasing population of *U · C · ADP* (Fig. 6b)— to prevent the substrate from aggregation. To maximize substrate folding, higher nucleotide exchange rate should accompany higher stimulated ATP hydrolysis rate (Fig. 6c, d, e).

**Figure 5.**
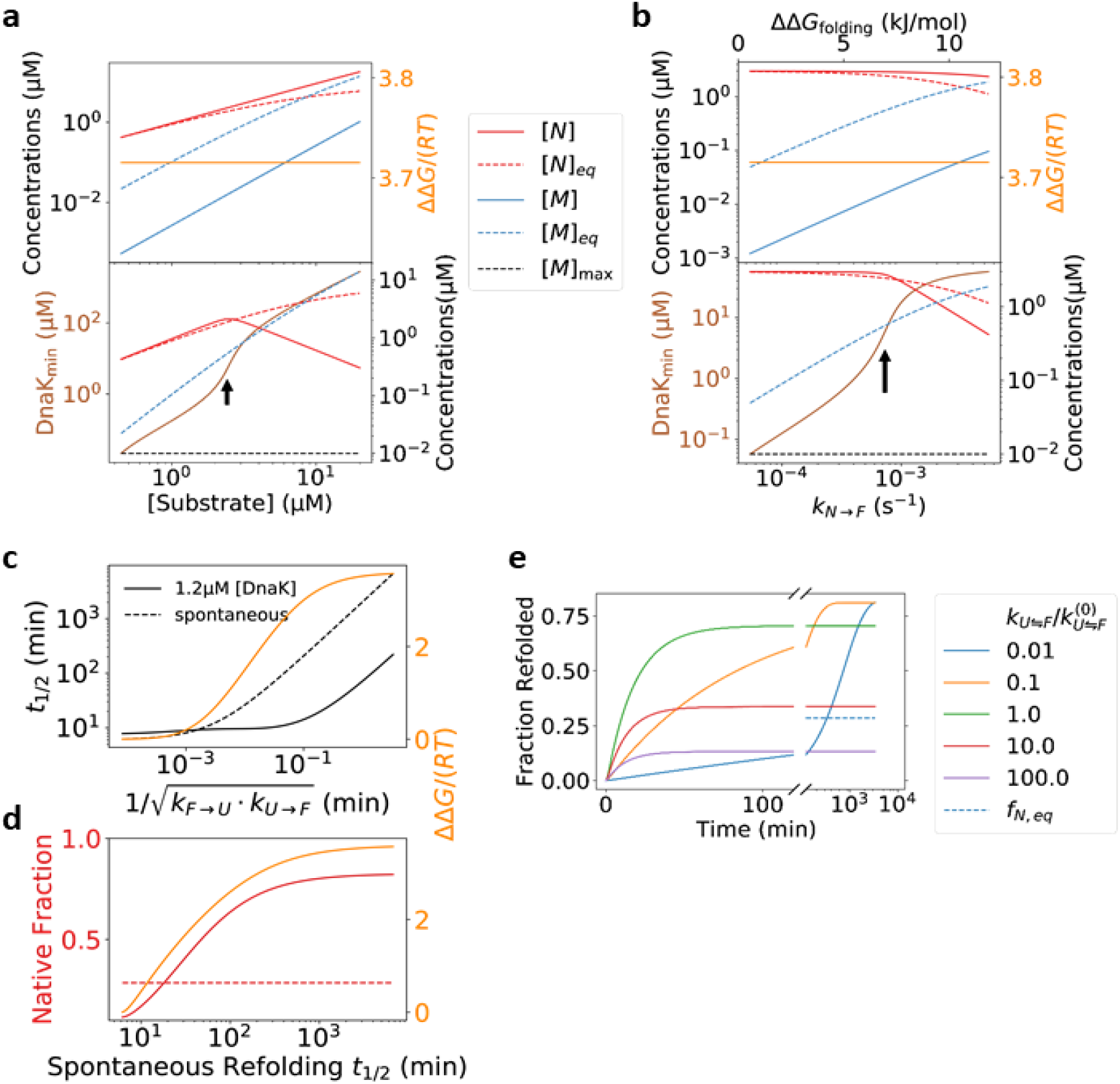
The capacity of the DnaK/DnaJ/GrpE chaperones to prevent aggregation and to elevate native protein concentrations. **a** The chaperones can prevent aggregation at increasing substrate concentrations. The rate of aggregation is taken to be proportional to the substrate concentration (see *Methods*). Top: the native and the misfolded steady state concentrations at increasing total LucDHis6 concentrations, with the DnaK concentration fixed at 1.2 μM. Bottom: the DnaK concentration required to maintain the misfolded protein concentration at or below [*M*]_max_ = 0.01 µM, as well as the steady state concentration of the native substrate at that DnaK concentration. **b** The chaperones can prevent aggregation at decreasing substrate stability. We vary the protein stability by changing the rate constant of conversion, *k*_*N*→*F*_, from the *N* state to the *F* state; the corresponding change in the folding free energy ΔΔ*G*_folding_ is indicated on the top axis. The native and the misfolded concentrations, as well as the DnaK concentration required to prevent aggregation, are shown as in panel a. **c** and **d** Hsp70 is more efficient at folding substrates with slower conversion between the *U* and the *F* states. Here, we take the kinetic parameters of luciferase folding at 25 °C, and simultaneously scale the forward and reverse rates of the reaction 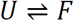 by the same factor, thus changing the kinetics without affecting the folding equilibrium. The times, *t*_1/*2*_, for the refolding of the misfolded substrate to reach half of the native fraction at equilibrium (spontaneous refolding) or the steady state (mediated by DnaK/DnaJ/GrpE), as well as the excess free energy (orange, right y-axis), are plotted against the hypothetic rates of conversion in c. The native fractions (red, left y-axis) and the excess free energy (orange, right y-axis) at the steady state are plotted against *t*_1/2_ of spontaneous refolding in d; the equilibrium native fraction is shown as the red dotted line. **e** The time courses of Hsp70-mediated refolding of the misfolded substrate at different hypothetical rates of conversion between *U* and *F* (keeping 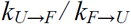 constant). Higher steady state native fractions are obtained at the price of longer refolding times.

**Figure 6.**
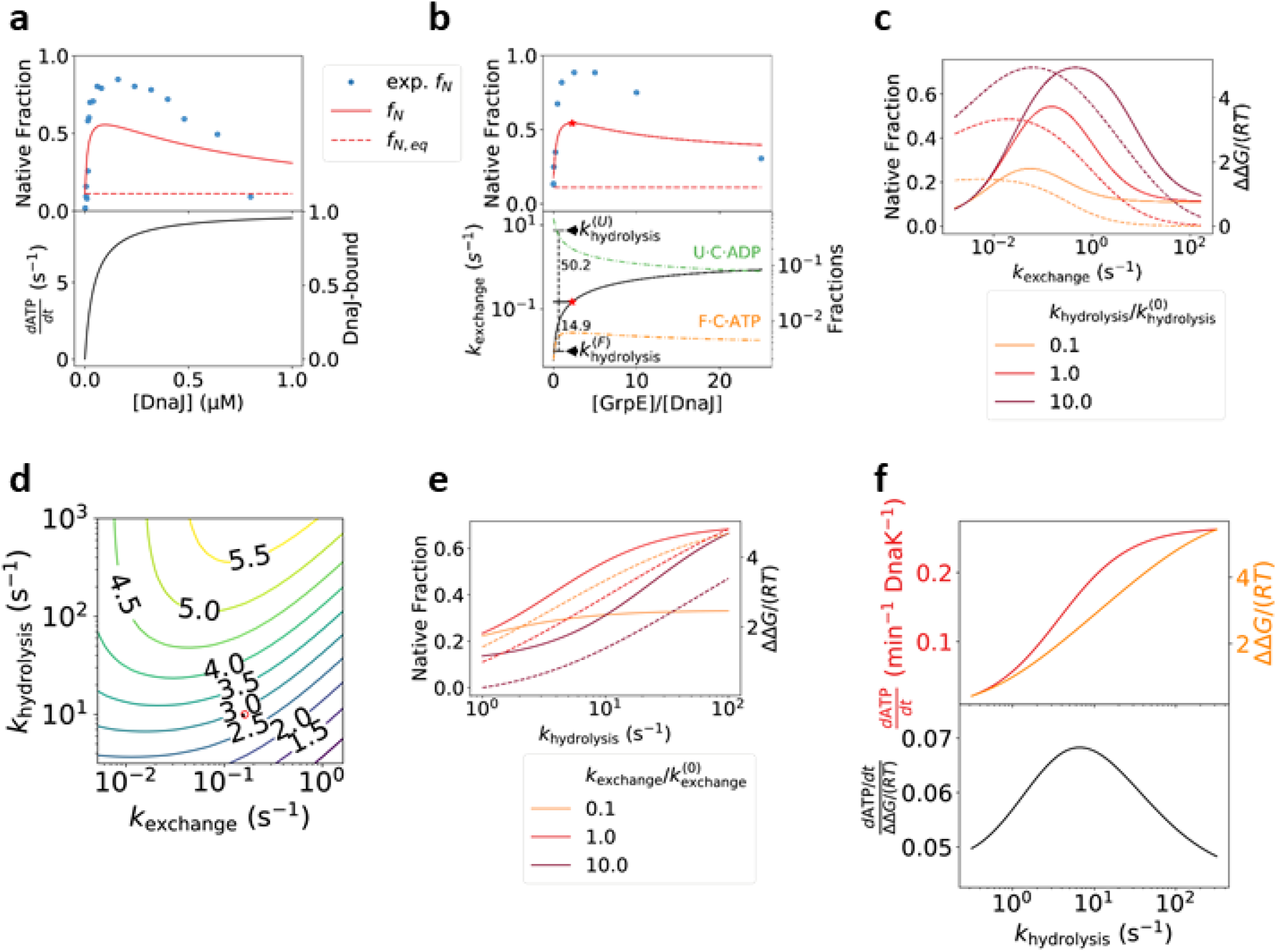
The rates of Hsp40-catalyzed ATP hydrolysis and NEF-catalyzed nucleotide exchange affect the efficiency of Hsp70-mediated folding. **a** Refolding of luciferase at various DnaJ concentrations. Top: the steady state native fractions predicted by our model, compared to the experimental data of refolding after 30 min at 30 °C ^15^. Bottom: the ATP hydrolysis rate (left y-axis) and the fraction of substrate bound to DnaJ (right y-axis). **b** Refolding of luciferase at various GrpE concentrations. Top: the steady state native fractions predicted by our model, compared to the experimental data of refolding after 2 hours at 30 °C ^27^. Bottom: the rate of nucleotide exchange at different GrpE concentrations (black line, left y-axis), and the populations of *F · C · ATP* and *U · C · ADP* (right y-axis). The optimal GrpE concentration is indicated by the red stars, and the in-plot numbers show the corresponding ratios of the nucleotide exchange rate to the ATP hydrolysis rates in the *U* and *F* states. In a and b, the rates of GrpE-catalyzed nucleotide exchange and DnaJ-catalyzed ATP hydrolysis are adjusted for the temperature of 30 °C (see *Methods* and Table 1). **c** Folding efficiency at different hypothetical rates of nucleotide exchange, for different values of the ATP hydrolysis rate in the *U* state. Native fractions (solid lines, left y-axis) are diminished at both low and high nucleotide exchange rates. At high rates of nucleotide exchange, the excess free energies (dashed lines, right y-axis) approach zero, indicating that Hsp70 can no longer drive protein folding. **d** The excess free energy as a function of the nucleotide exchange and the DnaJ-catalyzed ATP-hydrolysis rates. The rates used to model DnaK/DnaJ/GrpE-mediated folding at 30 °C are indicated by the red circle. **e** Folding efficiency increases with the DnaJ-catalyzed ATP hydrolysis rate, yielding higher native fractions (solid lines, left y-axis) and larger excessive free energies (dashed lines, right y-axis). **f** Higher ATP hydrolysis rate yields larger excess free energy (orange, right y-axis, top), at the price of higher rate of ATP consumption (red, left y-axis, top). The ratio of the two (bottom) changes only slightly.

**Table 3.** The conditions of the refolding experiments (the corresponding references are in superscripts). **A**. Spontaneous refolding of guanidinium chloride (GdmCl)-denatured luciferase. **B**-**D**. DnaK/DnaJ/GrpE-mediated refolding of GdmCl -denatured luciferase. **E**. DnaK/DnaJ/GrpE-mediated refolding of LucDHis6 denatured by 4 freeze-thaw cycles. LucDHis6 is an engineered luciferase variant where the last 62 COOH-terminal residues are replaced by SKLSYEQDGLHAGSPAALEHHHHHH-COOH.

|  | **A**<sup>4</sup> | **B**<sup>5</sup> | **C**<sup>15</sup> | **D**<sup>27</sup> | **E**<sup>29</sup> |
| --- | --- | --- | --- | --- | --- |
| [DnaK] (μM) | 0 | 1.25 | 0.8 | 0.8 | 0 to 6 |
| [DnaJ] (μM) | 0 | 0.25 | 0 to 0.8 | 0.16 | 0.3 |
| [GrpE] (μM) | 0 | 1.25 | 0.4 | 0 to 4 | 0.8 |
| [S] (μM) | 0.0324 | 0.25 | 0.08 | 0.08 | 3 |
| [ATP] (mM) | 0 | 1 | 5 | N/A | 5 |
| [ADP] (mM) | 0 | 0 | 0 | N/A | 0 |
| T (°C) | 20, 30 | 25 | 30 | 30 | 22 |

Our model predicts that Hsp70 chaperones with higher Hsp40-stimulated ATP hydrolysis rates can drive substrate folding to higher native fractions (Fig. 6e), at the cost of higher free energy expenditure (Fig. 6f). This result explains a previous experimental observation that a small molecule that enhances ATP hydrolysis by Hsp40-bound Hsp70 can induce higher yields of substrate folding^41^. Modulation of the ATP hydrolysis or the nucleotide exchange rates by small molecules may represent a therapeutic opportunity in the treatment of misfolding- or proteostasis-related diseases^42^.

## Discussion

Our model makes two distinct predictions that subject it to future experimental tests and possible falsification. First, it predicts that some thermodynamically unstable substrates depend on continuous Hsp70 assistance to maintain their native structures, and such a substrate in the steady state of Hsp70-mediated folding will gradually lose its native structure upon disruption of the chaperone system. Second, it predicts that an Hsp70 molecule bound to a substrate molecule will dissociate faster if subsequently the substrate molecule folds into the native state than if the substrate molecule misfolds, and such a difference will vanish in the absence of Hsp40. The second prediction may be tested by single molecule experiments^23,32^, if, for instance, separate fluorescence signals are available to detect Hsp70-substrate binding and substrate folding.

In support of the first prediction above, a recent experiment has demonstrated that luciferase at 37 °C can be kept active by the DnaK/DnaJ/GrpE chaperone system when there is sufficient ATP, but it rapidly loses its activity when ATP is depleted by the addition of apyrase^11^. Here, based on our model, we propose an alternative experiment, which avoids the complication that apyrase also affects the luciferase assay: Hsp70-mediated maintenance of luciferase activity may be disrupted by inhibiting the simultaneous binding of Hsp40 to Hsp70 and the substrate protein, which can be implemented, for example, by adding a J-domain (e.g., DnaJ with its CTD deleted) in excess to the chaperone system.

## Methods

### Model of Hsp70-mediated protein folding

We denote Hsp70 as *C*, Hsp40 (J protein) as *J*, and the NEF as *E*. [*Y*] denotes the solution concentration of the molecular species *Y*. There are four types of reactions explicitly considered in our model (Fig. 1b):

1) Hsp70 binding to the substrate.

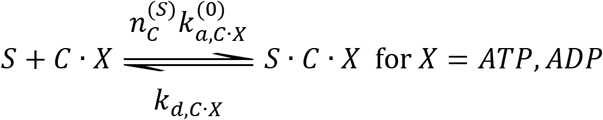

2) Conformational transitions of the substrate. An Hsp70-free substrate can adopt any of the four conformational states

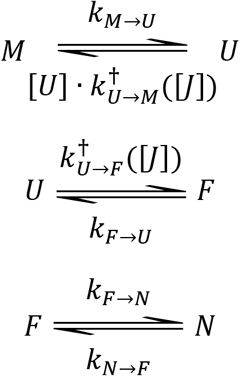

The chaperone-bound substrate can only be in and transition between the *U* and *F* states

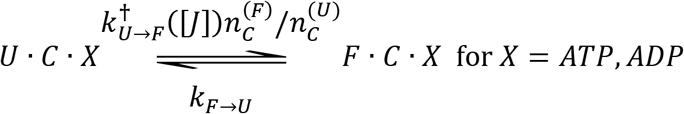

3) ATP hydrolysis.

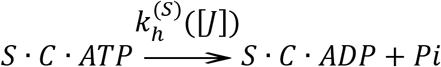

4) Nucleotide exchange.

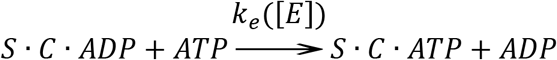

The details of the kinetic rates of the above reactions are described below.

### Hsp70 binding to the substrate

The association rate constant of Hsp70 binding to a substrate in state 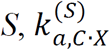, is determined by the number of accessible binding sites, 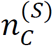, in that conformation: 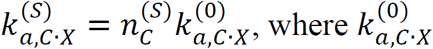 is the association rate constant of the chaperone binding to a fully accessible binding site, and *X = ATP, ADP.* The dissociation rate constant of Hsp70 from the substrate, *k*_*d,C*_._*x*_, does not depend on the substrate conformation, but depends on whether nucleotide *X = ATP or ADP* is bound. Experimentally, 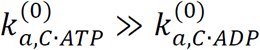 and 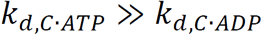. We assume 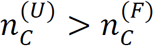.

### Conformational transitions of the substrate

The transition rates between conformations *S* and *S’* are different between a chaperone-free substrate 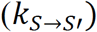 and a chaperone-bound substrate 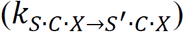 (Fig. 1b). The condition of thermodynamic cycle closure dictates that

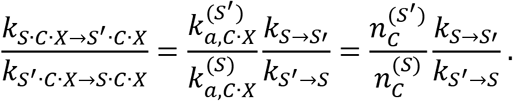

Because Hsp40 has different affinities for different substrate conformations, the transition rates between the conformations will depend on whether the substrate is bound to Hsp40. We treat the effects of Hsp40 on the reactions implicitly by making the affected rate constants dependent on the solution Hsp40 concentration [*J*] (see below).

For the transition 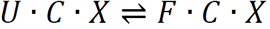, we assume that the bound chaperone does not hinder the substrate to go from the *F* state to the *U* state, because a binding site available in the *F* state is most likely also available in the *U* state (based on the assumption 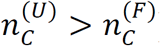. Thus we take 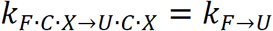. It follows from thermodynamic cycle closure that the rate of the reverse transition—we use the superscript dagger to indicate that they are influenced by the presence of Hsp40—is

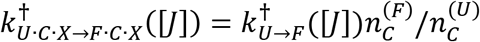

We take the rate of aggregation to be proportional to the substrate concentration:

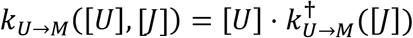

The rates 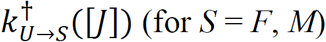 (for *S = F, M*) depend on the affinities of Hsp40 for the substrate in different conformational states. For simplicity, we assume that Hsp40 only binds to the substrate in the *U* state, and consequently only the Hsp40-free substrate can change from conformation *U to F* or *M.* The corresponding transition rates are

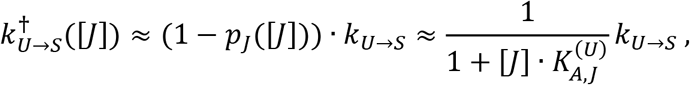

where *k*_*U*→ *S*_ is the rate of transition *U* → *S* (for *S = F, M*) for an Hsp40-free substrate, *p*_*J*_ is the probability that the substrate is Hsp40-bound, and 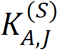 is the binding constant of Hsp40 for the conformational state *S.*

### Hsp40-substrate binding

To keep our model simple, we do not explicitly consider the kinetics of binding and unbinding between Hsp40 and the substrate, and make the approximation that they are always at equilibrium. The key assumption of our model is that the fold-competent conformation *F* is much less accessible to Hsp40 than the aggregation-prone conformation *U,* i.e., 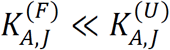. To reduce the number of unknown parameters, we take 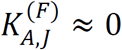, i.e., the binding of Hsp40 to the fold-competent conformation is negligible, as in our derivation of the transition rates above. We also neglect subtleties such as that the J domain may bind with different affinities to Hsp70 in the ATP- and ADP-states^43^. The above approximations may contribute to quantitative differences between the predictions of our model and the experimental observations, particularly in predicting how folding changes with Hsp40 concentrations. The binding and unbinding of Hsp40 to the substrate and to Hsp70 can be explicitly included in our model at the cost of greater complexity and additional fitting parameters, but our simplified treatment above is adequate for the key results in this work.

### ATP hydrolysis

The Hsp40-stimulated Hsp70 ATP hydrolysis rate, 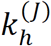, can be orders-of-magnitude higher than the unstimulated basal rate 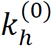. The ATP hydrolysis rate of Hsp70 bound to the substrate in the *F* state is simply 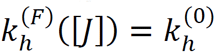, following our approximation that no Hsp40 binds to the substrate in the *F* state. When Hsp70 is bound to the substrate in the *U* state, its average rate of ATP hydrolysis, given the solution Hsp40 concentration, [*J*], can be approximated by

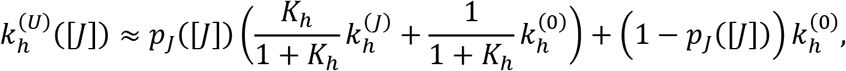

where 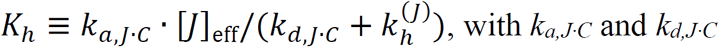 being the association and dissociation rates of J domain binding to Hsp70, and [*J*]eff being the effective concentration of a substrate-bound Hsp40 molecule around an Hsp70 molecule bound to the same substrate molecule. The first term on the right hand side is the steady state rate of catalysis, weighted by the probability *p*_*J*_ that an Hsp40 is bound to the substrate and thus present to catalyze the hydrolysis. Here we assume a high ATP concentration such that ATP binding to Hsp70 is fast compared to other steps in ATP hydrolysis.

### NEF-catalyzed nucleotide exchange

Because of the high concentration of ATP in cells and in the refolding experiments, we treat this reaction as irreversible. The reaction proceeds in three steps: 1) dissociation of ADP, 2) binding of ATP, and 3) conformational change of Hsp70 from the closed conformation in the ADP-state to the open conformation in the ATP-state. The conformational equilibrium between the open and closed conformations may be influenced by Hsp40 binding to Hsp70^44^, but this effect is not considered in our model for simplicity and lack of experimental parameters. In absence of the nucleotide exchange factor, the rate limiting step in the reaction is the dissociation of ADP, with the rate constant *k*_*d,ADP*_, whereas when catalyzed by the NEF, the rate limiting step is the conformational change, with rate constant *k*_*C*_ ^27,45^. The overall rate of reaction at a given NEF concentration, [*E*], is then approximately

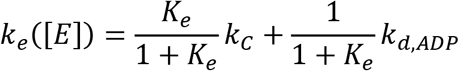

where 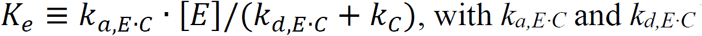 being the association and dissociation rates of NEF binding to Hsp70. The temperature dependence of *k*_*C*_ for DnaK has been determined to satisfy the Arrhenius equation: 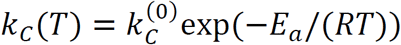, where *R* = 8.314 J/mol/K is the gas constant.

### Solving the kinetic equations

To simplify the calculations of refolding kinetics, we make the approximation that the solution concentrations of Hsp70, Hsp40, and NEF remain constant throughout the refolding process, which is true if they are in large excess of the substrate-bound chaperone, cochaperone, and NEF. Under this approximation, refolding kinetics is described by a set of linear ordinary differential equations, which are solved by the technique of eigenvalue decomposition of the rate matrix. This simplification allows quick and robust fitting of the folding kinetic parameters to the experimental refolding data. The steady state calculations do not use this approximation.

### Proof that without cochaperones Hsp70 cannot alter the population ratio between the native and the misfolded states

Consider a hypothetical, single-conformation substrate *s* and a reference reaction cycle of chaperone binding 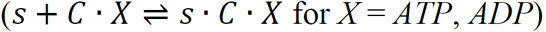, ATP hydrolysis 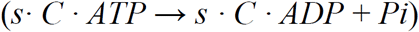, and nucleotide exchange (*s · C · ADP + ATP* → *s · C · ATP + ADP*), where the ATP-bound and ADP-bound chaperones bind to *s* with the rate constants 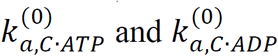, respectively. Let 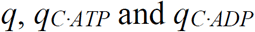 be the fractions of the hypothetical substrate that are chaperone-free (*s*), bound to an ATP-bound chaperone (*s · C · ATP*), and bound to an ADP-bound chaperone (*s · C · ADP*), respectively, at the steady state of this reaction cycle. For the real substrate in Hsp70-mediated folding, let *f*_*S*_ and *f*_*S·C·X*_ (*X = ATP, ADP*) be the fractions of the free and the *C · X*-bound substrates in conformation *S,* and let *f*_*S,eq*_ be the fraction of the substrate in conformation *S* at the folding equilibrium in the absence of the chaperones. It can be verified that

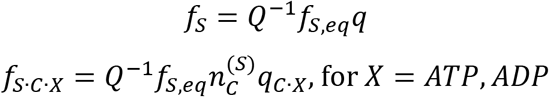

where

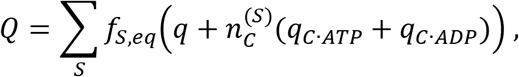

are the steady state solutions to the kinetic equations, given the condition of thermodynamic cycle closure. Thus the ratio 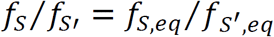 is not altered by the chaperone, despite the free energy expenditure of ATP hydrolysis. This holds true for all numbers of intermediate states, so long as the ATP hydrolysis and the nucleotide exchange rates of the chaperone do not depend on the conformational state of the bound substrate.

## Author Contributions

HX designed and performed the research and wrote the paper.

## References

1. Bukau, B., Weissman, J. & Horwich, A. Molecular chaperones and protein quality control. Cell 125, 443–451 (2006).

2. Hartl, F. U., Bracher, A. & Hayer-Hartl, M. Molecular chaperones in protein folding and proteostasis. Nature 475, 324–332 (2011).

3. Balchin, D., Hayer-Hartl, M. & Hartl, F. U. In vivo aspects of protein folding and quality control. Science 353, aac4354 (2016).

4. Herbst, R., Schäfer, U. & Seckler, R. Equilibrium intermediates in the reversible unfolding of firefly (Photinus pyralis) luciferase. J. Biol. Chem. 272, 7099–7105 (1997).

5. Szabo, A., Langer, T., Schroder, H., Flanagan, J., Bukau, B. & Hartl, F. U. The ATP hydrolysis-dependent reaction cycle of the Escherichia coli Hsp70 system DnaK, DnaJ, and GrpE. Proc. Natl. Acad. Sci. 91, 10345–10349 (1994).

6. Mayer, M. P., Schröder, H., Rüdiger, S., Paal, K., Laufen, T. & Bukau, B. Multistep mechanism of substrate binding determines chaperone activity of Hsp70. Nat. Struct. Biol. 7, 586–593 (2000).

7. Mayer, M. P. Hsp70 chaperone dynamics and molecular mechanism. Trends in Biochemical Sciences 38, 507–514 (2013).

8. Barducci, A. & De Los Rios, P. Non-equilibrium conformational dynamics in the function of molecular chaperones. Current Opinion in Structural Biology 30, 161–169 (2015).

9. Goloubinoff, P. & Rios, P. D. L. The mechanism of Hsp70 chaperones: (entropic) pulling the models together. Trends in Biochemical Sciences 32, 372–380 (2007).

10. Santra, M., Farrell, D. W. & Dill, K. A. Bacterial proteostasis balances energy and chaperone utilization efficiently. Proc. Natl. Acad. Sci. 114, E2654–E2661 (2017).

11. Goloubinoff, P., Sassi, A. S., Fauvet, B., Barducci, A. & Rios, P. D. L. Chaperones convert the energy from ATP into the nonequilibrium stabilization of native proteins. Nat. Chem. Biol. 14, 388–395 (2018).

12. Kampinga, H. H. & Craig, E. A. The HSP70 chaperone machinery: J proteins as drivers of functional specificity. Nature Reviews Molecular Cell Biology 11, 579–592 (2010).

13. Yang, J., Nune, M., Zong, Y., Zhou, L. & Liu, Q. Close and allosteric opening of the polypeptide-binding site in a human Hsp70 chaperone BiP. Structure 23, 2191–2203 (2015).

14. Kityk, R., Kopp, J., Sinning, I. & Mayer, M. P. Structure and dynamics of the ATP-bound open conformation of Hsp70 chaperones. Mol. Cell 48, 863–874 (2012).

15. Laufen, T., Mayer, M. P., Beisel, C., Klostermeier, D., Mogk, A., Reinstein, J. & Bukau, B. Mechanism of regulation of Hsp70 chaperones by DnaJ cochaperones. Proc. Natl. Acad. Sci. 96, 5452–5457 (1999).

16. Alderson, T. R. R., Kim, J. H. H. & Markley, J. L. L. Dynamical structures of Hsp70 and Hsp70-Hsp40 complexes. Structure 24, 1014–1030 (2016).

17. Wittung-Stafshede, P., Guidry, J., Horne, B. E. & Landry, S. J. The J-Domain of Hsp40 couples ATP hydrolysis to substrate capture in Hsp70. Biochemistry 42, 4937–4944 (2003).

18. Szabo, A., Korszun, R., Hartl, F. U. & Flanagan, J. A zinc finger-like domain of the molecular chaperone DnaJ is involved in binding to denatured protein substrates. EMBO J. 15, 408–417 (1996).

19. Lu, Z. & Cyr, D. M. The conserved carboxyl terminus and Zinc finger-like domain of the co-chaperone Ydj1 assist Hsp70 in protein folding. J. Biol. Chem. 273, 5970–5978 (1998).

20. Perales-Calvo, J., Muga, A. & Moro, F. Role of DnaJ G/F-rich domain in conformational recognition and binding of protein substrates. J. Biol. Chem. 285, 34231–34239 (2010).

21. Rüdiger, S., Schneider-Mergener, J. & Bukau, B. Its substrate specificity characterizes the DnaJ co-chaperone as a scanning factor for the DnaK chaperone. EMBO J. 20, 1042–1050 (2001).

22. Sekhar, A., Velyvis, A., Zoltsman, G., Rosenzweig, R., Bouvignies, G. & Kay, L. E. Conserved conformational selection mechanism of Hsp70 chaperone-substrate interactions. Elife 7, e32764 (2018).

23. Mashaghi, A., Bezrukavnikov, S., Minde, D. P., Wentink, A. S., Kityk, R., Zachmann-Brand, B., Mayer, M. P., Kramer, G., Bukau, B. & Tans, S. J. Alternative modes of client binding enable functional plasticity of Hsp70. Nature 539, 448–451 (2016).

24. Ahmad, A., Bhattacharya, A., McDonald, R. A., Cordes, M., Ellington, B., Bertelsen, E. B. & Zuiderweg, E. R. P. Heat shock protein 70 kDa chaperone/DnaJ cochaperone complex employs an unusual dynamic interface. Proc. Natl. Acad. Sci. 108, 18966–18971 (2011).

25. Han, W. & Christen, P. Mechanism of the targeting action of DnaJ in the DnaK molecular chaperone system. J. Biol. Chem. 278, 19038–19043 (2003).

26. Johnson, J. L. & Craig, E. A. An Essential Role for the Substrate-Binding Region of Hsp40s in Saccharomyces cerevisiae. J. Cell Biol. 152, 851–856 (2001).

27. Packschies, L., Theyssen, H., Buchberger, A., Bukau, B., Goody, R. S. & Reinstein, J. GrpE accelerates nucleotide exchange of the molecular chaperone DnaK with an associative displacement mechanism. Biochemistry 36, 3417–3422 (1997).

28. Brehmer, D., Rüdiger, S., Gässler, C. S., Klostermeier, D., Packschies, L., Reinstein, J., Mayer, M. P. & Bukau, B. Tuning of chaperone activity of Hsp70 proteins by modulation of nucleotide exchange. Nat. Struct. Biol. 8, 427–432 (2001).

29. Sharma, S. K., Rios, P. D. L., Christen, P., Lustig, A. & Goloubinoff, P. The kinetic parameters and energy cost of the Hsp70 chaperone as a polypeptide unfoldase. Nat. Chem. Biol. 6, 914–920 (2010).

30. Rios, P. D. L. & Barducci, A. Hsp70 chaperones are non-equilibrium machines that achieve ultra-affinity by energy consumption. Elife 3, e02218 (2014).

31. Astumian, R. D. Microscopic reversibility as the organizing principle of molecular machines. Nat. Nanotechnol. 7, 684–688 (2012).

32. Nunes, J. M., Mayer-Hartl, M., Hartl, F. U. & Müller, D. J. Action of the Hsp70 chaperone system observed with single proteins. Nat. Commun. 6, 6307 (2015).

33. Hu, B., Mayer, M. P. & Tomita, M. Modeling Hsp70-mediated protein folding. Biophys. J. 91, 496–507 (2006).

34. Diamant, S., Peres Ben-Zvi, A., Bukau, B. & Goloubinoff, P. Size-dependent disaggregation of stable protein aggregates by the DnaK chaperone machinery. J. Biol. Chem. 275, 21107–21113 (2000).

35. Diamant, S. & Goloubinoff, P. Temperature-controlled activity of DnaK-DnaJ-GrpE chaperones: Protein-folding arrest and recovery during and after heat shock depends on the substrate protein and the GrpE concentration. Biochemistry 37, 9688–9694 (1998).

36. Goloubinoff, P., Mogk, A., Zvi, A. P. Ben, Tomoyasu, T. & Bukau, B. Sequential mechanism of solubilization and refolding of stable protein aggregates by a bichaperone network. Proc. Natl. Acad. Sci. 96, 13732–13737 (1999).

37. Calloni, G., Chen, T., Schermann, S. M., Chang, H., Genevaux, P., Agostini, F., Tartaglia, G. G., Hayer-Hartl, M. & Hartl, F. U. DnaK functions as a central hub in the E. coli chaperone network. Cell Rep. 1, 251–264 (2012).

38. Aguilar-Rodríguez, J., Sabater-Muñoz, B., Montagud-Martínez, R., Berlanga, V., Alvarez Ponce, D., Wagner, A. & Fares, M. A. The molecular chaperone DnaK is a source of mutational robustness. Genome Biol. Evol. 8, 2979–2991 (2016).

39. Murugan, A., Huse, D. A. & Leibler, S. Speed, dissipation, and error in kinetic proofreading. Proc. Natl. Acad. Sci. 109, 12034–12039 (2012).

40. Hopfield, J. J. Kinetic Proofreading: A New Mechanism for Reducing Errors in Biosynthetic Processes Requiring High Specificity. Proc. Natl. Acad. Sci. 71, 4135–4139 (1974).

41. Wisén, S., Bertelsen, E. B., Thompson, A. D., Patury, S., Ung, P., Chang, L., Evans, C. G., Walter, G. M., Wipf, P., Carlson, H. A., Brodsky, J. L., Zuiderweg, E. R. P. & Gestwicki, J. E. Binding of a small molecule at a protein-protein interface regulates the chaperone activity of Hsp70-Hsp40. ACS Chem. Biol. 5, 611–622 (2010).

42. Labbadia, J. & Morimoto, R. I. The Biology of proteostasis in aging and disease. Annu. Rev. Biochem. 84, 435–464 (2015).

43. Kim, J. H., Alderson, T. R., Frederick, R. O. & Markley, J. L. Nucleotide-dependent interactions within a specialized Hsp70/Hsp40 complex involved in Fe–S cluster biogenesis. J. Am. Chem. Soc. 136, 11586–11589 (2014).

44. Yang, J., Zong, Y., Su, J., Li, H., Zhu, H., Columbus, L., Zhou, L. & Liu, Q. Conformation transitions of the polypeptide-binding pocket support an active substrate release from Hsp70s. Nat. Commun. 8, s41467–17 (2017).

45. Slepenkov, S. V & Witt, S. N. Kinetics of the reactions of the Escherichia coli molecular chaperone DnaK with ATP: evidence that a three-step reaction precedes ATP hydrolysis. Biochemistry 37, 1015–1024 (1998).

46. Suh, W.-C., Burkholder, W. F., Lu, C. Z., Zhao, X., Gottesman, M. E. & Gross, C. A. Interaction of the Hsp70 molecular chaperone, DnaK, with its cochaperone DnaJ. Proc. Natl. Acad. Sci. 95, 15223–15228 (1998).

47. Russell, R., Jordan, R. & McMacken, R. Kinetic characterization of the ATPase cycle of the DnaK molecular chaperone. Biochemistry 37, 596–607 (1998).

48. McCarty, J. S., Buchberger, A., Reinstein, J. & Bukau, B. The role of ATP in the functional cycle of the DnaK chaperone system. J. Mol. Biol. 249, 126–137 (1995).

